# CpG Island Definition and Methylation Mapping of the T2T-YAO Genome

**DOI:** 10.1101/2023.12.02.568720

**Authors:** Ming Xiao, Rui Wei, Jun Yu, Chujie Gao, Fengyi Yang, Le Zhang

## Abstract

Precisely defining and mapping all cytosine positions and their clusters, known as CpG islands (CGIs), as well as their methylation status are pivotal for genome-wide epigenetic studies, especially when population-centric reference genomes are ready for timely application. Here we first align the two high-quality reference genomes, T2T-YAO and T2T-CHM13, from different ethnic backgrounds in a base-by-base fashion and compute their genome-wide density-defined and position-defined CGIs. Second, mapping some representative genome-wide methylation data from selected organs onto the two genomes, we find that there are about 4.7–5.8% sequence divergency of variable categories depending on quality cutoffs. Genes among the divergent sequences are mostly associated with neurological functions. Moreover, CGIs associated with the divergent sequences are significantly different with respect to CpG density and observed CpG/expected CpG (O/E) ratio between the two genomes. Finally, we find that the T2T-YAO genome not only has a greater CpG site coverage than that of the T2T-CHM13 genome when whole-genome bisulfite sequencing (WGBS) data from the European and American populations are mapped to each reference, but also show more hyper-methylated CpG sites as compared to the T2T-CHM13 genome. Our study suggests that future genome-wide epigenetic studies of the Chinese populations rely on both acquisition of high-quality methylation data and subsequent precision CGI mapping based on the Chinese T2T reference.

## Introduction

Position-sensitive and co-methylated CpG islands (CGIs) represent one of the several key epigenetic mechanisms for chromosome integrity and gene expression regulation [1, 2]. To define highly variable CGIs based on both position and density genome-wide and to map their methylation status under different physiological conditions and in different cell types are prerequisite in most epigenomic studies [3–7]. Human reference genomes, especially those tailored to specific populations, provide single-base-resolution coordinates for genome-wide genetic and epigenetic mapping with ultimate precision [8]. The release of the telomere-to-telomere CHM13 (T2T-CHM13) genome in 2022 has provided such a benchmark by filling-up sequence gaps that contribute nearly 8% increase of the previous human genome assembly [9]. Therefore, it is of essence to compare it to the more recent human genome assembly of the Chinese population, the T2T-YAO genome [10], and to annotate all CGIs for both assemblies.

Here, we simply raise three basic questions. First, are there significant differences in CGIs and their methylation between the two reference genomes? And if so, what are the relevant genes and their functions diverged between them? Second, does the T2T YAO genome provide additional information or serve more appropriately as one of the reference genomes for the Chinese population? Both are of importance for subsequent genome-wide epigenetic studies and building methylation databases specifically for the Chinese populations under complex physiological and pathological conditions. We, therefore, chose to accurately predict genome-wide CGIs for both reference genomes and validate their applications in methylation studies by testing them on a limited number of datasets due to data availability and quality.

A genome-wide comparison of two reference genomes has been done recently between the classical GRCH38 and T2T-CHM13 genomes and the authors discovered nearly 200 million base pairs (Mbp) unique to the T2T-CHM13 genome[9]. This result supports our effort in mapping the T2T-YAO and T2T-CHM13 genomes to explore their divergent sequences, and determining whether the divergent sequences are associated with specific biological functions became our first scientific question. We have described previously a genome-wide inclusive CGI definition, *i.e.*, a distance-based clustering method yielding short CpG cluster, known as position-defined CGIs show potential gene regulatory functions and are closely correlated with gene expression specificity [2, 11, 12]. Therefore, whether there was any specificity between T2T-YAO and T2T-CHM13 according to CGI features, became our second scientific question. It becomes feasible now to carry out a comparative analysis on the T2T-YAO and T2T-CHM13 genomes on CGI methylation profiles using public WGBS data [2, 8, 11, 12], and this is what we address as the third scientific question.

To this end, we align the two genomes, compute both density-defined and position defined CGIs, separately, and map some genome-wide methylation WGBS data from representative organs to analyze differentially methylated CGIs between the two genomes. Our major findings are as follows: (1) There is an approximately 4.7–5.8% difference between the two genomes, which is composed of divergent sequence associated genes correlated closely with neurological functions based on GO enrichment analysis [13–18]. (2) CGIs associated with the divergent sequences are significantly different with respect to CpG density and O/E ratio [11, 19]. (3) Not only do WGBS data mapped to the T2T-YAO genome have a greater CpG site coverage than those of the T2T-CHM13 genome, but also the T2T-YAO genome shows more hyper methylated CpG sites than T2T-CHM13 does. Our study demonstrates statistically significant differences between T2T-YAO and T2T-CHM13 in genome sequence and epigenetic landmarks[20–22], such as CGI and methylation patterns, suggesting that the establishment of CGI and methylation profiles based on the Chinese T2T reference genome is crucial for subsequent genome-wide epigenetic studies based on Chinese populations.

## Results

In order to address the three scientific questions (**Figure 1)**, we analyzed the two genomic references, their CGI distribution, and genome-wide CGI-associated methylation.

**Figure 1.**
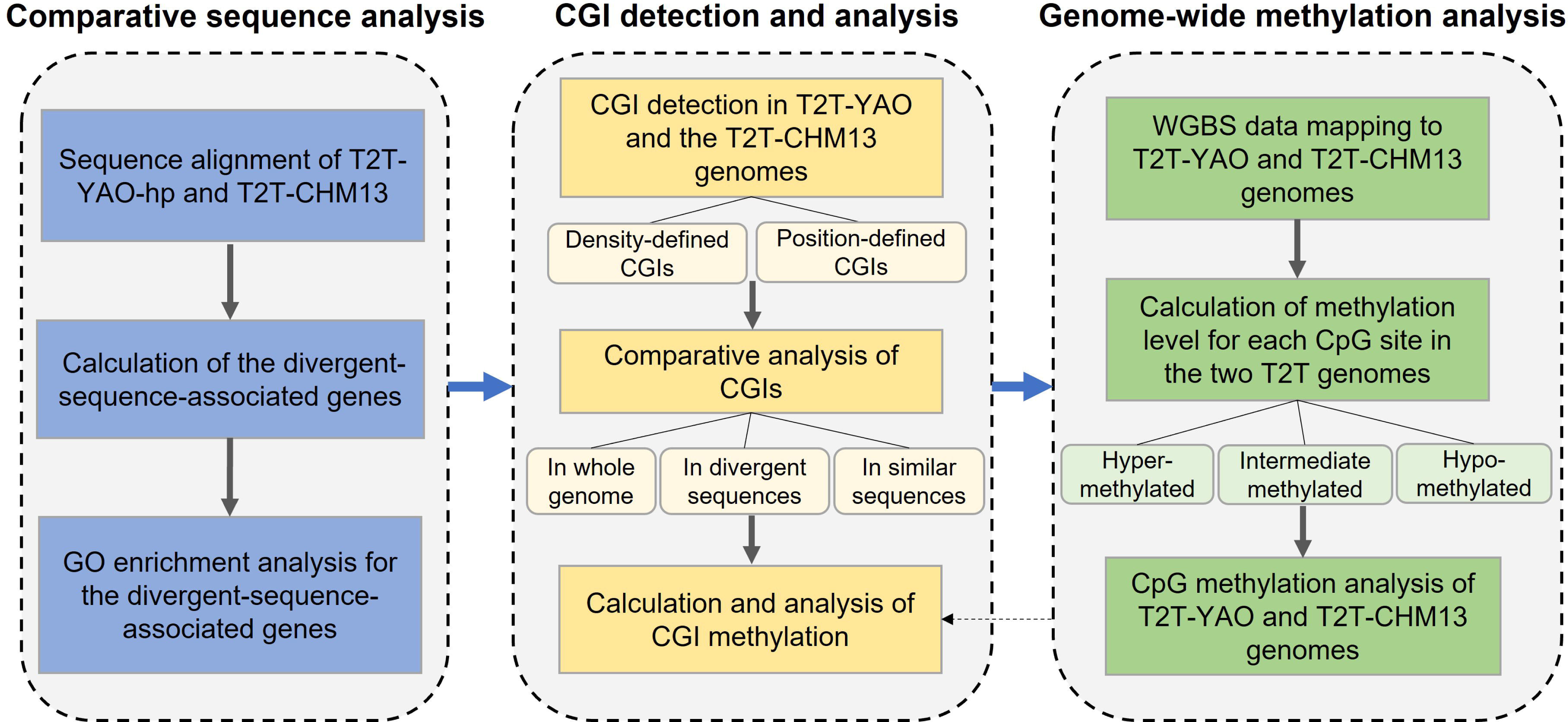
Workflow of the study. GO, Gene Ontology; T2T, telomere-to-telomere; WGBS, whole-genome bisulfite sequencing; CGI, CpG island.

### Comparative sequence analysis

To address our first question, we aligned the T2T-YAO-hp and T2T-CHM13 genomes and functionally categorized the divergently enriched genes from the T2T-YAO-hp genome (*-hp* stands for the haploid sequence from the offspring of the triad family T2T-YAO assemblies; T2T-YAO-hp and T2T-YAO used here are equivalent unless otherwise specified).

#### Sequence differences between T2T-YAO-hp and T2T-CHM13

First, from the sequence alignment (the alignment results are detailed in Supplementary **Table S1 and S2**), as shown in **Table 1**, the T2T-CHM13 genome has approximately 54.6 Mbp more sequence as compared to the T2T-YAO-hp genome. The uniquely divergent sequences account for 5.751% and 4.665% of the full length T2T-CHM13 and T2T-YAO-hp genomes, respectively. Therefore, the difference between the two genomes is approximately 4.7%–5.8%. Second, using NGenomeSyn [23], we created a collinearity map of the two genomes and observed that the two sequences are very similar both in identity and chromosome length (**Figure 2)** in general [24]. The divergent sequences between the two genomes are mainly concentrated in specific regions of each chromosome (Figure 2B). For instance, some of the divergent sequences of chromosomes 15 and 22 are enriched at the their left telomeric ends, whereas others, such as chromosome 9, have those located near the centromeres. The examples of some longest diverge sequence segments include 3.31 × 10^7^ bp *vs*. 1.69 × 10^7^ bp of chromosome 9 and 2.15 × 10^7^ bp *vs.* 1.31 × 10^7^ bp of chromosome Y; both are longer in the T2T-CHM13 genome (Figure 2C).

**Figure 2.**
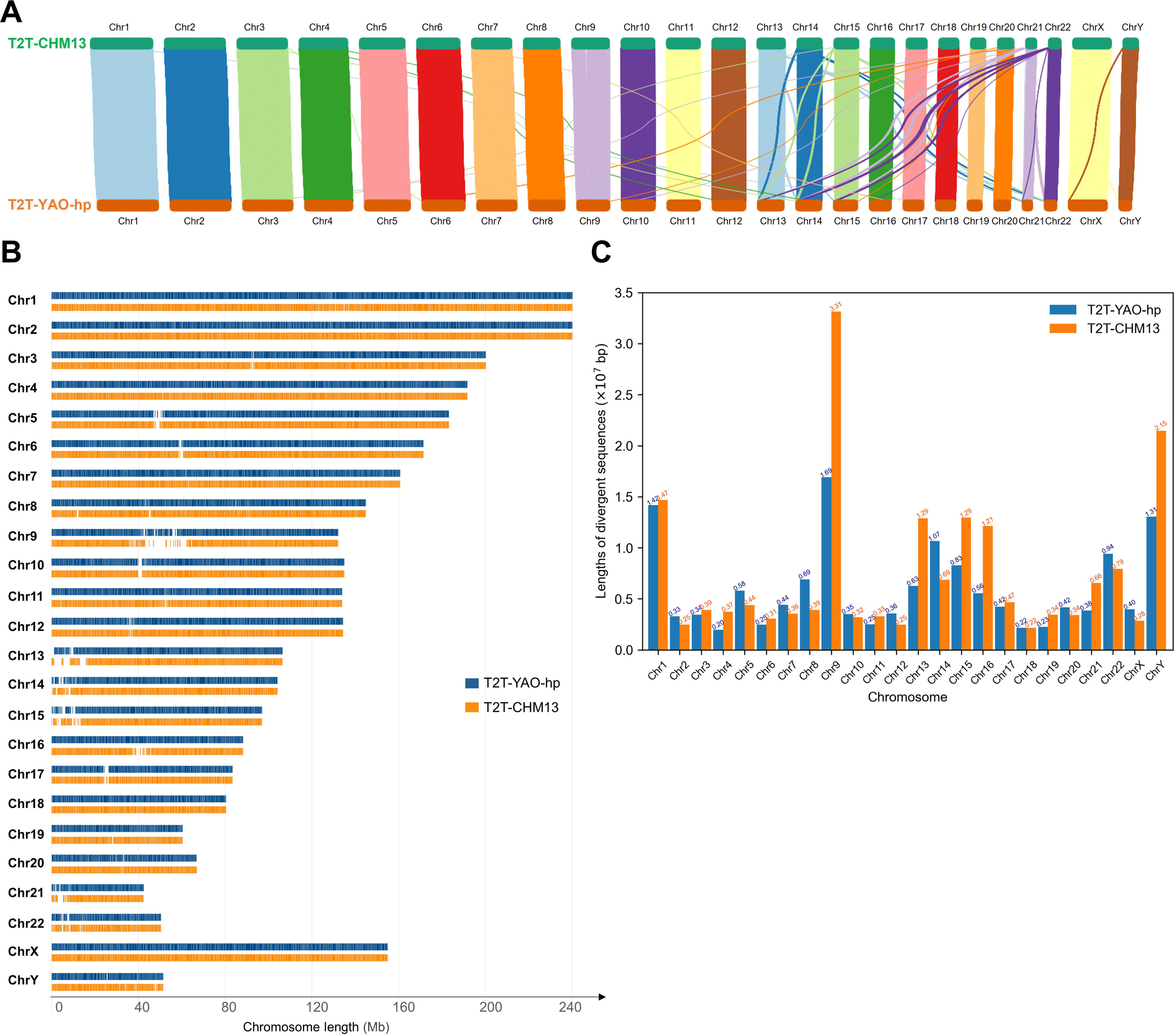
Sequence alignment between the T2T-CHM13 and T2T-YAO-hp genomes. **A.** A collinearity map of the genome alignment. Colored lines link the high-identity or collinear sequence segments of the two compared genomes and the line thickness as a similarity measure. **B.** Visualization of sequence comparison across chromosomes. Blue and orange highlight similar sequences as vertical bars of the T2T-CHM13 and T2T-YAO-hp genomes, respectively. Blank bars represent divergent sequences of the two genomes. **C.** Divergent sequence length statistics of the two genomes. Blue and orange bars represent the divergent sequence lengths of the T2T-CHM13 and T2T-YAO-hp genomes in different chromosomes, respectively. Chr, chromosome.

**Table 1.**
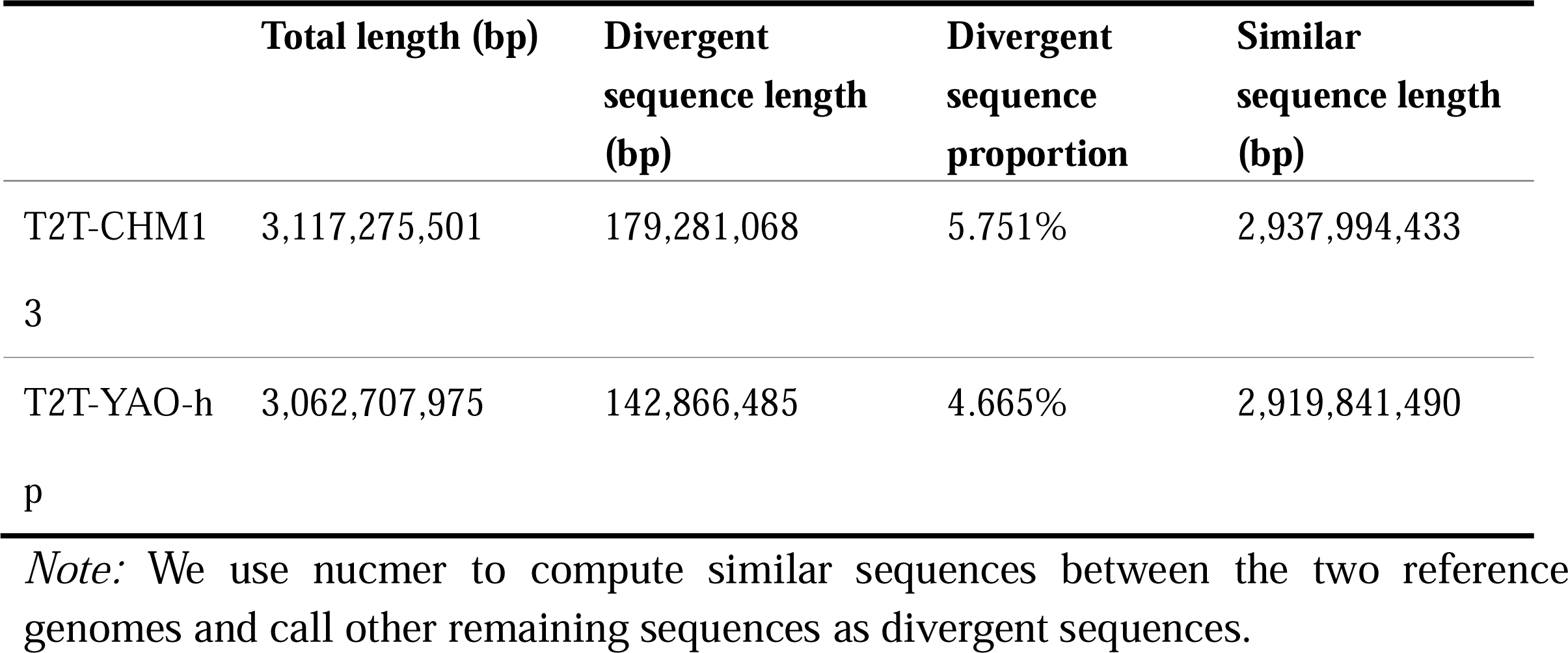
Sequence differences between the T2T-CHM13 and T2T-YAO-hp genomes.

#### Divergent-sequence-associated genes

We categorized the divergent-sequence-associated genes of the two genomes (**Figure 3A** and Supplementary **Table S3)** and show that most of them are protein-coding genes, accounted for 44.26%, and the rest include IncRNAs that are 25.93% of the total gene count. Our GO enrichment analysis for the protein-coding genes shows that the most significant terms are “homophilic cell adhesion via plasma membrane adhesion molecules”, “ion channel complex”, and “gated channel activity” in the biological process (BP), cellular component (CC), and molecular function (MF) categories [25], respectively, and that most of these GO terms are associated with certain functions of the nervous system (**Figure 3B)**.

**Figure 3.**
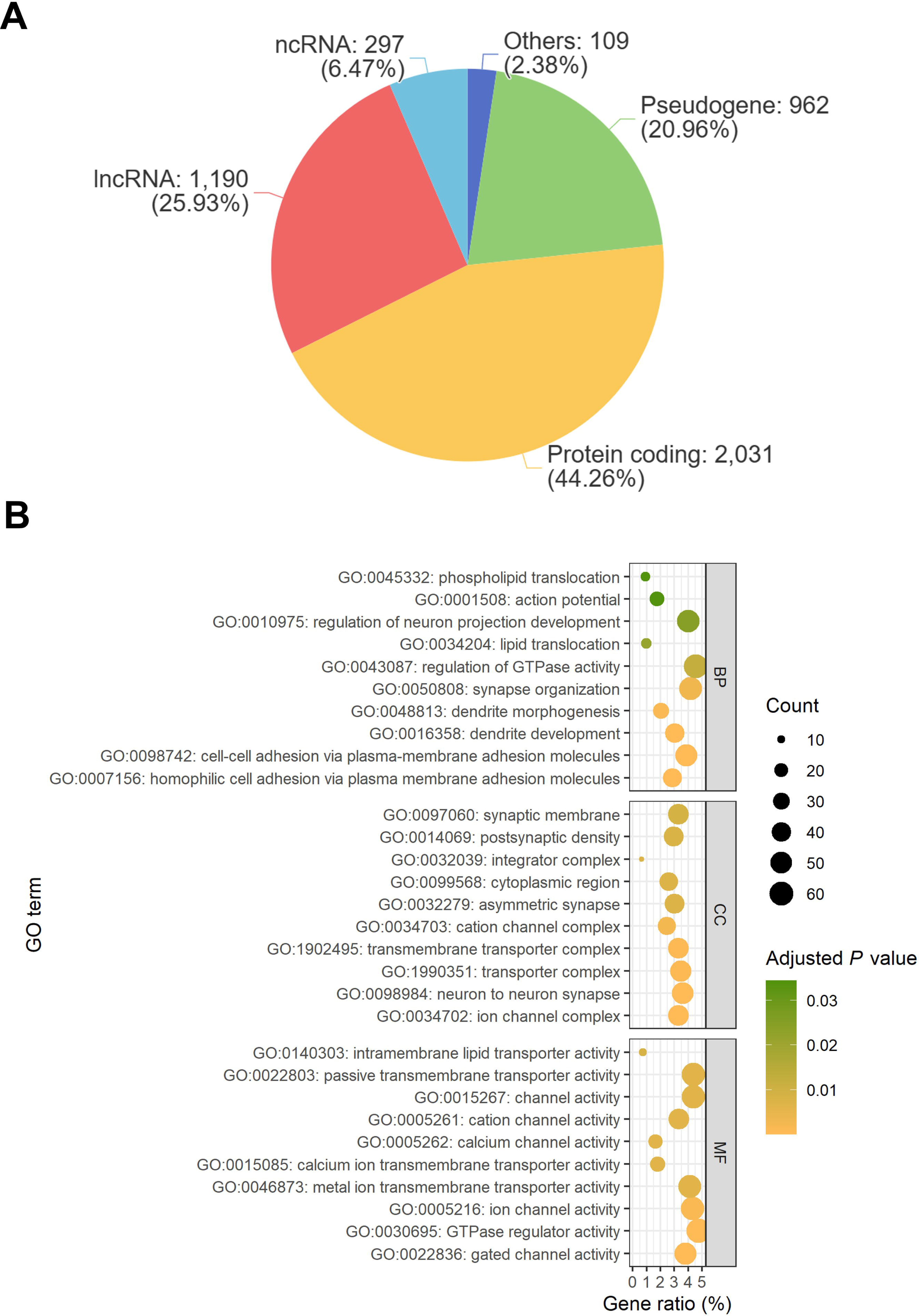
GO enrichment of divergent-sequence-associated genes. **A.** Proportion of the types of divergent-sequence-associated genes in these two reference genomes. **B.** GO enrichment analysis of divergent sequence related protein-coding genes. Size and color of the circle represent the number of enriched genes and the statistical significance value (adjusted *P* value) [22, 41] of the current GO term, respectively. ncRNA, non-coding RNA; lncRNA, long non-coding RNA; BP, biological process; MF, molecular function; CC, cellular component.

### CGI prediction and analysis

#### CGI prediction results

We employed the rule-based method [12, 19] and distance-based clustering method [12, 19] to predict the number of CGIs and the average CGI length for the two genomes (**Table 2** and Supplementary **Table S4)**, and show the distribution of CGIs on each of the T2T-YAO-hp and T2T-CHM13 chromosomes (Supplementary **Figure S1** compares). On one hand, the number of density-defined CGIs and the average length predicted for the T2T-YAO-hp genome are slightly smaller (84,471 bp *vs.* 87,788 bp and shorter (391 bp *vs.* 407 bp) than those of the T2T-CHM13 genome, respectively. On the other, the number of position-defined CGIs predicted for the T2T-YAO-hp genome is greater than those of the T2T-CHM13 genome under the default parameter d=50 bp (222,421 bp *vs.* 220,736 bp). In addition, the CGI number predicted for the T2T-YAO-hp genome is less than those of the T2T-CHM13 genome regardless if their density-defined CGIs or position-defined CGIs, but the average length of the position-defined CGIs predicted by the T2T-YAO-hp genome is collectively longer than those of the T2T-CHM13 genome (*e.g.*, 60 bp *vs.* 58 bp with d = 25 bp) under the parameters d = 25 bp and 12 bp. We also predicted all CGIs related to divergent sequences of the two genomes (Supplementary **Table S5)**.

**Table 2.**
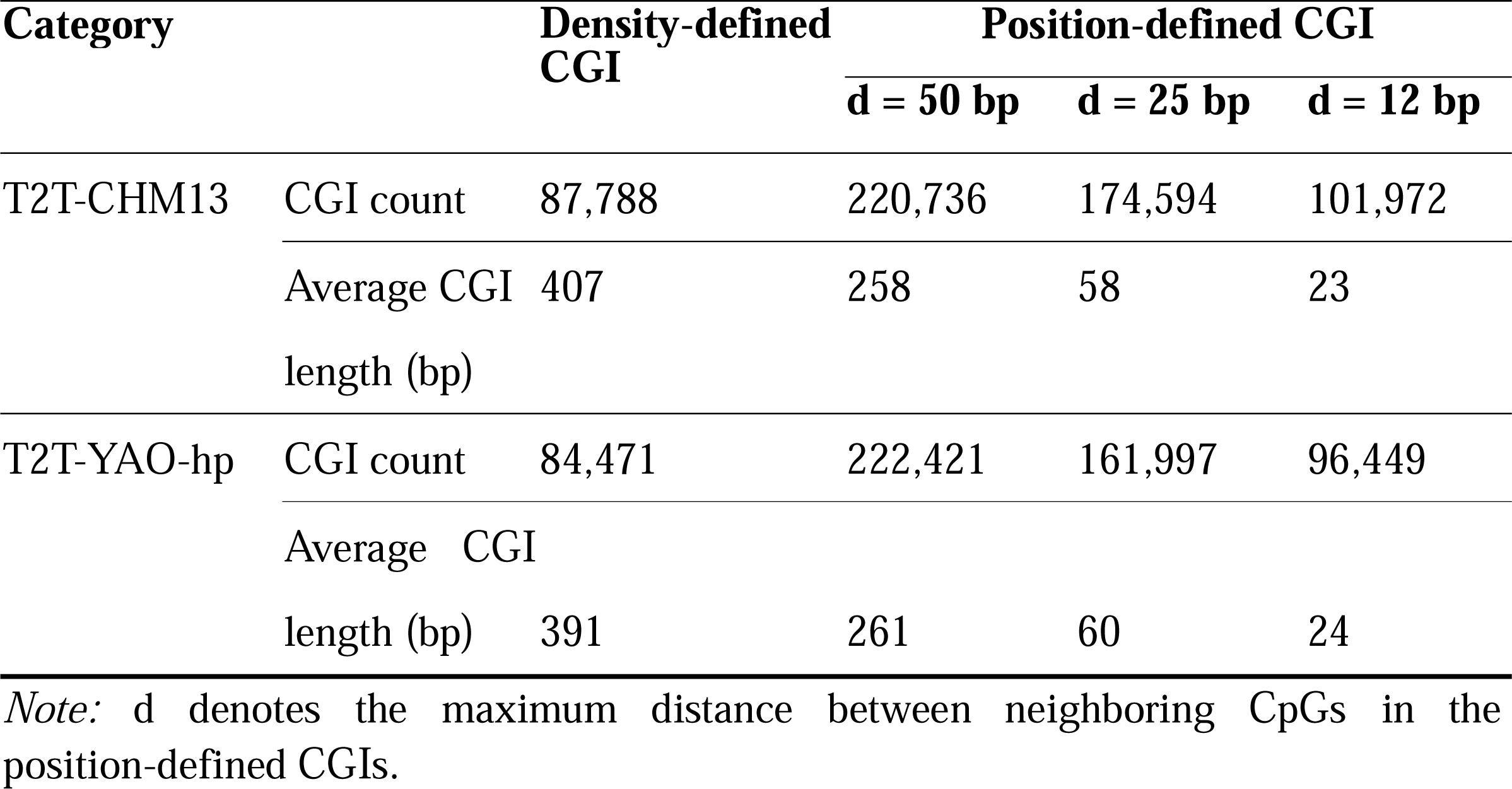
CGIs Statistics of the T2T-CHM13 and T2T-YAO-hp genomes.

To study the relationship between position-defined CGIs and density-defined CGIs for the same genome, we investigated sequences that intersect the two groups (for the position-defined CGIs, d = 25 bp) for both genomes **(Figure 4)**. First, the total length of density-defined CGIs is much longer, approximately three times that of position-defined CGIs for each genome *(e.g.*, (24,991,625 + 8,062,542)/ (8,062,542 + 1,735,619) = 3.37 for the T2T-YAO-hp genome). Second, although most of the position-defined CGIs overlap with the density-defined CGIs, there is still a fraction (16.5% and 17.7%, respectively) that is not predicted by the rule-based CGI predicting method.

**Figure 4.**
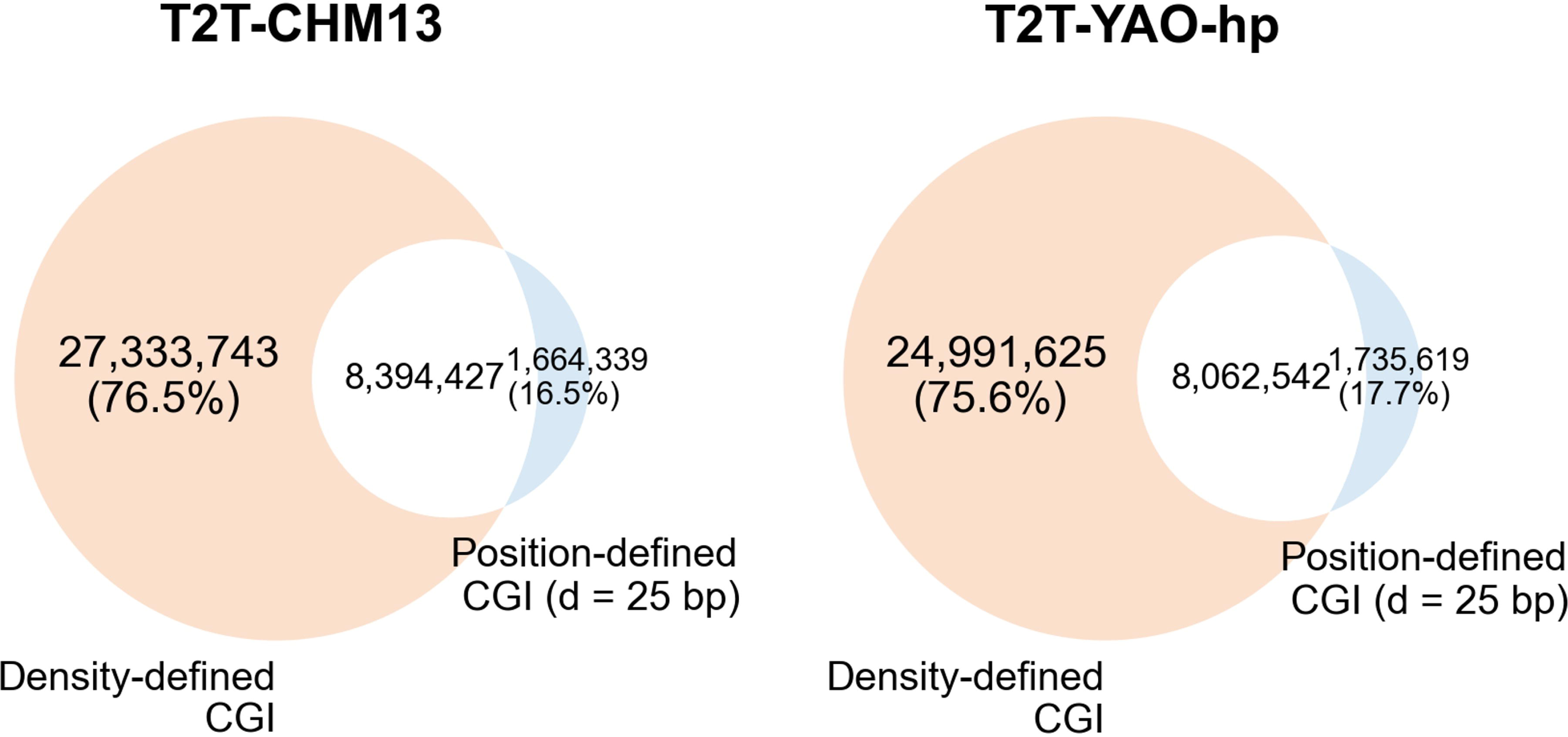
The sequence intersecting the density-defined and position-defined CGIs (d = 25 bp). The white part represents the sequence length intersecting position-defined and density-defined CGIs. The orange and blue parts represent the sequence length unique to the density-defined and position-defined CGIs, respectively.

#### CGI feature comparison

First, to compare the differences in CGIs between the two genomes, we calculated four commonly used CGI features, *i.e.*, CGI length, GC content, CpG density, and CGI O/E ratio [2, 26] (**Figure 5**) and show the feature distribution of genome-wide and divergent-sequence-associated position-defined CGIs, including d = 12 bp, 50 bp (Supplementary **Figure S2)**. Other than the grouping of density-defined or position-defined CGIs, there is no obvious difference among the four commonly used CGI features between the two genomes based on all CGI counts (Figure 5A–5D). For example, in both genomes, the peak lengths of the density-defined CGIs and the position-defined CGIs are approximately 210 bp and 30–50 bp, respectively (Figure 5A).

**Figure 5.**
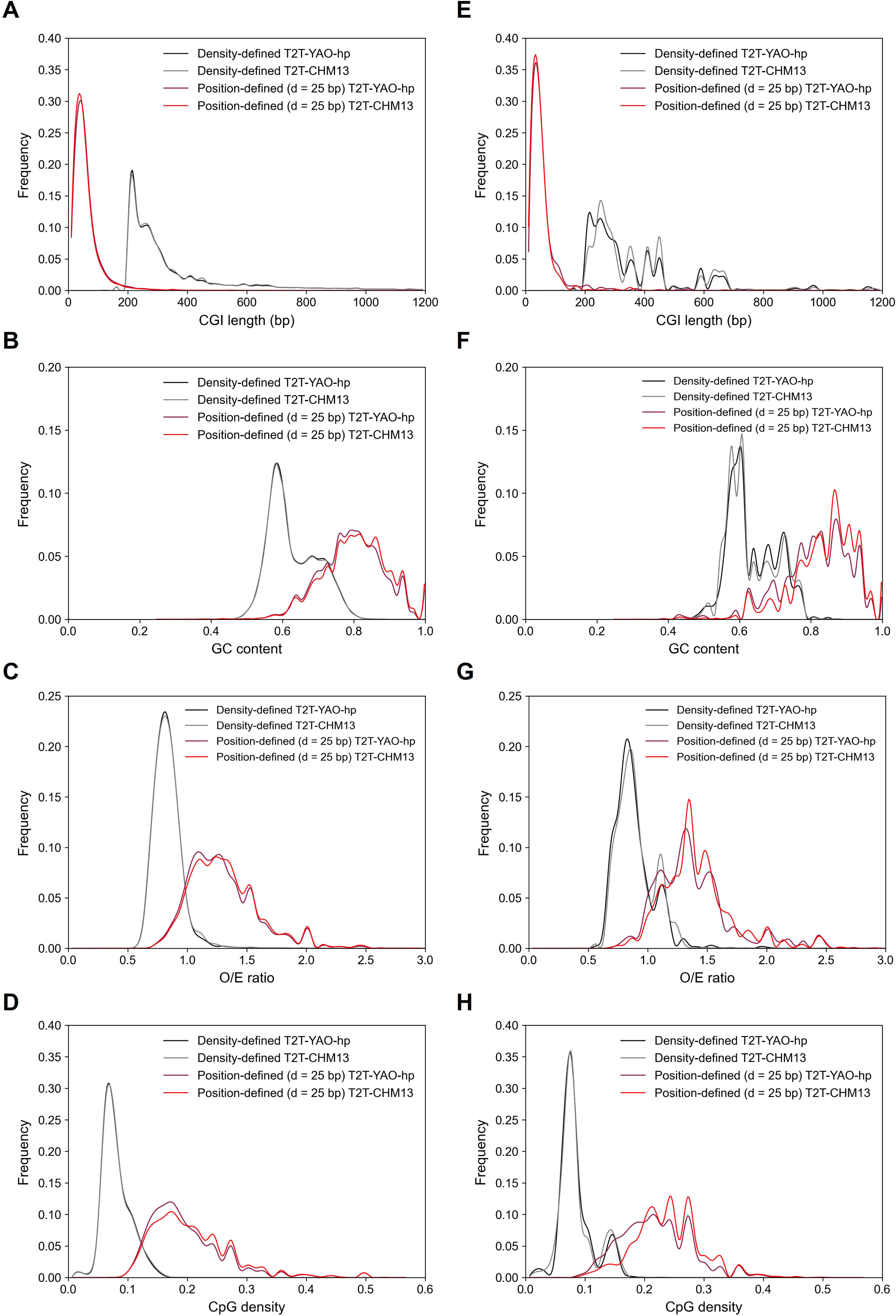
Comparison of CGI features between the T2T-CHM13 and T2T-YAO-hp genomes. **A.** Length of all CGIs; **B.** GC content of all CGIs; **C.** O/E ratio of all CGIs; **D.** CpG density of all CGIs; **E.** Length of divergent-sequence-associated CGIs; **F.** GC content of divergent-sequence-associated CGIs; **G.** O/E ratio of divergent-sequence-associated CGIs; **H.** CpG density of divergent-sequence-associated CGIs. O/E, observed CpG/expected CpG.

Second, there is a greater difference in the four commonly used CGI features of the divergent sequence CGIs between the two genomes than those of all CGI counts (Figure 5E–5H). For instance, the density-defined CGIs of the T2T-CHM13 genome have two peaks in high GC content regions (from 50% to 60%), whereas the density-defined CGIs of the T2T-YAO-hp genome have a single peak in the same regions. In addition, the position-defined CGIs in the divergent sequences, except for a small difference in the distribution of CGI lengths, have significantly different distributions for GC content, O/E ratio, and CpG density (Figure 5E–5H, Supplementary **Figure S2**, and Supplementary **Table S6**).

Third, for all CGIs or divergent-sequence-associated CGIs, the distance-based clustering method is more likely to identify CGIs with shorter lengths and greater GC content, O/E ratio, and CpG density than the rule-based CGI predicting method (Figure 5).

### Methylation analysis and comparison

To address our third question, we compared not only the number of cytosine sites (C sites, Supplementary **Figure S3**) and methylation levels of all CpG sites, but also methylation levels of genome-wide CGIs for the two genomes.

#### CpG methylation levels

We classified CpG methylation levels of similar (Supplementary **Figure S4**) and divergent sequences between the two genomes (**Figure 6**). In addition, based on the WGBS data of two germline tissues, the ovary and testis, we also determined methylation profiles of whole genomes and divergent sequences for the two genomes (Supplementary **Figure S5).** The number of CpG sites of the divergent sequences failed to be mapped based on the same WGBS methylation data is much lower in the T2T-YAO-hp genome than in the T2T-CHM13 genome in both germline tissues. For example, the number of CpG sites in the divergent sequences of the ovary failed to be mapped to T2T-YAO-hp is 2.16 × 10^6^ bp, whereas that in the T2T-CHM13 genome is 3.49 × 10^6^ bp. Inversely, the number of CpG sites in the divergent sequences mapped for T2T-YAO-hp is greater than that for T2T-CHM13 at all three methylation levels. In terms of quantity, the most notable difference is what found in the hypermethylated sequences (2.8 × 10^5^ *vs*. 2.2 × 10^5^ for ovary; 3.1 × 10^5^ *vs.* 2.5 × 10^5^ for testis).

**Figure 6.**
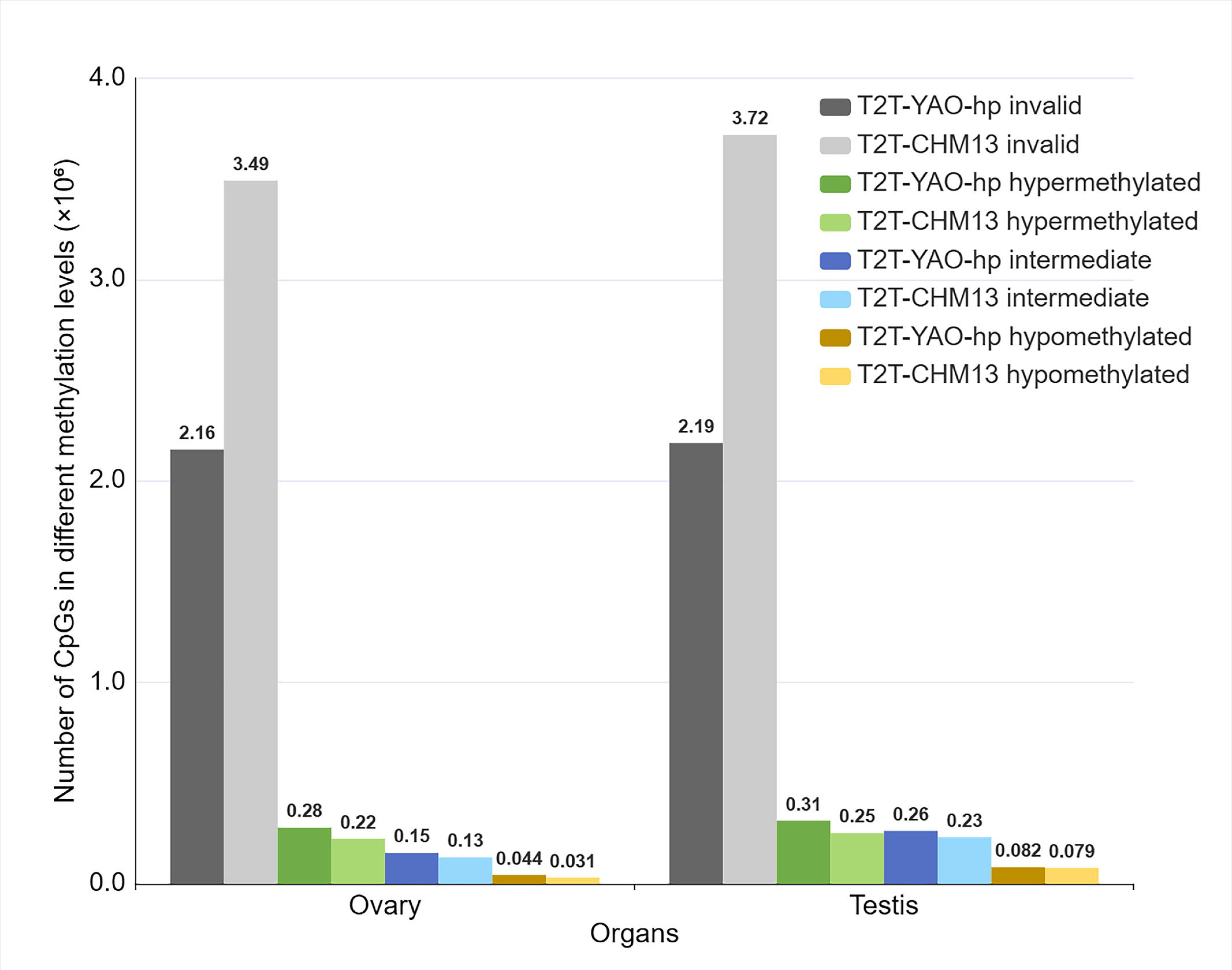
CpG methylation level of divergent sequences between the T2T-CHM13 and T2T-YAO-hp genomes. Green, blue, and orange highlight the number of hyper-methylated, intermediate methylated, and hypo-methylated, respectively. Gray highlights the number of CpG sites that failed in mapping with WGBS data. Dark and light colors are used to distinguish the T2T-YAO-hp and T2T-CHM13 genomes, respectively.

#### CGI methylation levels

We used Equation 6 to calculate the methylation level of all density-defined and position-defined CGIs for the two reference genomes (**Figure 7** and Supplementary **Figure S6**). First, after both definitions, the proportions of CGI-invalid data (WGBS data failed to be mapped to any CGIs) are higher in the T2T-CHM13 than in T2T-YAO-hp genomes (Figure 7); there are 10.46% *vs*. 7.27% for the density-defined CGIs and 23.63% *vs.* 14.87% for position-defined CGIs not mapped in the ovary for the two genomes (Figure 7A). Second, the proportions of hyper-methylated, intermediate methylated, and hypo-methylated of CGIs, predicted by both definitions, are slightly greater for the T2T-YAO-hp genome than those of the T2T-CHM13 genome; the proportions of hyper-methylated, intermediate methylated, and hypo-methylated for the T2T-YAO-hp genome predicted as position-defined CGIs are 19.73%, 13.29%, and 53%, respectively, as opposed to those of the T2T-CHM13 genome, 17.32%, 11.77%, and 48.02% (Figure 7B). Third, the density-defined CGIs appear to have the greatest proportion of hypermethylation for both genomes, whereas position-defined CGIs have a greater proportion of hypomethylation (Figure 7). The density-defined CGIs of both genomes have higher proportions of hyper-methylated CGIs (50.02% *vs.* 48.40% for the T2T-YAO and T2T-CHM13 genomes, respectively) than those of intermediate methylated and hypo-methylated CGIs, whereas the position-defined CGIs have the greatest proportion of hypomethylation (53.65% *vs.* 48.57% for the T2T-YAO and T2T-CHM13 genomes, respectively).

**Figure 7.**
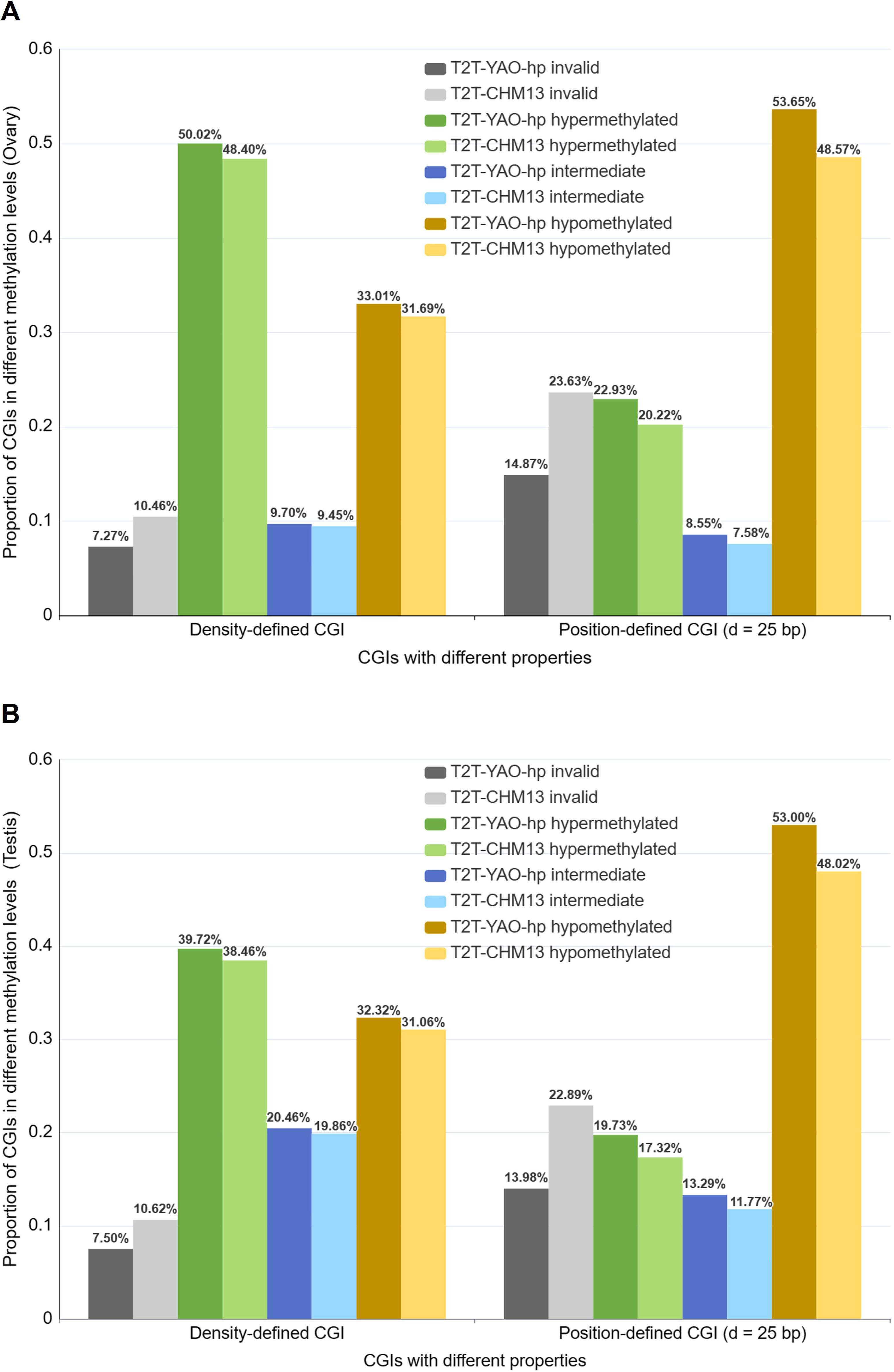
CGI methylation levels of the two genomes. **A.** CGI methylation levels annotated by using the ovary data. **B.** CGI methylation levels annotated by using the testis data. Green, blue, and orange highlight the proportion of CGIs that are hyper-methylated, intermediate methylated, and hypo-methylated, respectively. Gray highlights the proportion that failed from all CGIs mapping, denoted as *invalid*. Darker and lighter colors are used to distinguish the T2T-YAO-hp and T2T-CHM13 genome, respectively.

## Discussion and conclusion

The aim of this study is to emphasize merits and specific applications of a population-centric reference genome by comparing the sequence divergence, as well as various epigenetic parameters, between the T2T-YAO and T2T-CHM genomes. We have further provided the evidence suggesting the necessity of using T2T-YAO as a reference genome for future genome-wide genetic and epigenetic studies in Chinese populations. We have started with the two high-quality genome sequences to compute the genome-wide cytosine and CpG sites and further categorized them into two groups of CGIs, position-defined and density-defined. After the three-round stratification, we have presented the divergent-sequence-associated genes and concluded that these genes are mostly neurological in function. In addition, we have mapped the methylation levels of CpGs and CGIs for the two reference genomes using selected public WGBS data.

The overall sequence divergence between the two reference genomes are significant and interesting, albeit only 4.7–5.8% (minimal alignment length of 5,000bp and identity cutoff of 90%). Such a loosely-defined difference, generally speaking, is mostly population-associated variations between the northern Chinese and European populations [9, 10]. Future studies will have to expand empirical data into within-population variations based on big data for their detailed categorization of functional relevance. The current divergent sequences appear to non-uniformly spread out into certain chromosomal regions within a few chromosomes and their specific regions, such as the cases involving chromosomes 9 and Y; such a distribution also suggests population-associated selection and recombination. Furthermore, the GO enrichment of divergent-sequence-associated genes specific to the T2T-YAO genome indicates a strong preference for specific molecular mechanisms related to neuronal cells and synapses, providing clues for future in-depth investigation.

Our genome-wide CGI study between the two genomes indicates that two-fold CGI stratifications are necessary for precision methylation mapping. It is important to start methylation mapping from all potential methylation sites that are ensured only by T2T population-specific genomes. The two CGI definitions and their grouping schemes, such as the distance (12 bp, 25 bp, and 50 bp) and the density (high, moderate, and low), provide matrices for pattern and rule recognitions, which are used to identify genes and other genomic elements and features for further functional scrutinization. In addition, after the application of the two CGI definitions, there is still a significant fraction of the C and CpG sites to be further annotated, leveraging on the complete genomes, as some of these sites are actually methylated.

From the methylation mapping of the two datasets representing the two germline tissues, we have learnt several lessons. First, we expected more CpG sites to be mapped in the T2T-CHM13 reference from the methylation data of the European origins since its DNA is from a single homozygous complete hydatidiform mole (46, XX) and the Y sequence is from another assembly of similar origin. However, to our surprise, the number of CpG methylation sites to the divergent sequence associated to the T2T-YAO genome is greater than that of the T2T-CHM13 reference. Second, the CpG sites that failed to be mapped by the gender-specific methylation data are also biased toward the same direction, *i.e.*, there are more to be mapped in the T2T-YAO reference. These subjective mapping results lead to an across-the-board conclusion that methylation patterns are reference sensitive regardless the methylation levels. Third, we have also shown apparent differences of CGI methylation between the two CGI definitions and the rate varies among different methylation levels.

After all, there are still several questions that need further exploration. First, the success of large-scale genome-wide epigenomic studies rely on hign-quality and population-specific reference genomes and there are more much more references to be constructed for human methylation atlas of different populations. Second, novel and more thorough CpG and CGI definitions are needed for understanding the relatedness of data to be mapped and their references. Since artificial intelligence (AI) has been gradually used in genome-wide methylation-associated studies [27, 28], determining how to use AI techniques [29–32] such as neural networks to investigate the relationship between sequence features, CGI, methylation, and expression specificity of the human genome has become a future research direction. Finally, genome-wide CGI prediction and methylation analysis are not only data intensive but also time-consuming. However, there is currently no such web service that collects and manages CGI and methylation annotation data for the Chinese T2T genome. Therefore, we will build a web service for online computation, data sharing and visualization based on high-performance computing[33, 34], such as GPU acceleration and distributed computing, to provide a data foundation and an online platform for further study.

## Materials and methods

### Data source

We downloaded the human GRCh38 and T2T-CHM13 (v2.0) genome sequences from the NCBI databases [35] and the Chinese T2T-YAO-hp genome sequence from the BIGD (BIG Data Center, Beijing) databases [10]. We used CGI data of the T2T-CHM13 genome, which were predicted based on the Gardiner-Garden and Frommer (GGF) [36] standard in UCSC [37]. We downloaded the WGBS data for two representative tissues from the ENCODE database [38], which are ovary (ENCSR417YFD) from a European donor and testis (ENCSR806NNG) from an American individual.

### Methods

The workflow of this study in three main aspects is described in Figure 1, and our key algorithms used are described as follows.

### Comparative sequence analysis

#### Statistics of all cytosine C sites

We counted and compared the number and distribution of cytosine sites (C sites) that can be methylated among the T2T-YAO-hp, T2T-CHM13, and GRCh38 genomes. Since most DNA methylation sites found in mammals involve CpG dinucleotides [39], we categorized C sites into three different contexts: CpG, CHG, and CHH, where H represents A, C or T.

#### Genome alignment and analysis

We used nucmer, a sequence alignment algorithm of the MUMmer system [40], to carry out genome-wide alignment and analysis for the T2T-YAO-hp and T2T-CHM13 genomes (**Figure S7)**. We set the key parameters for nucmer to calculate similar and divergent sequences as follows: minimum alignment length is 5,000 bp, minimum alignment similarity is 90%. The similar sequences are those regions identified by nucmer as aligning between the two genomes. Conversely, divergent sequences are those regions that do not meet these above-mentioned alignment criteria, thus representing unique sequences present in each genome. The GO enrichment of divergent-sequence-associated genes were analyzed by using the clusterProfiler algorithm [41].

#### CGI definition and analysis

Our previous studies denoted the CGIs predicted by the classical rule-based method [36] as density-defined CGIs, and the CGIs predicted by the distance-based clustering method [26] as position-defined CGIs, which are able to localize more short CGIs of potential gene regulatory functions and are closely correlated with gene expression specificity [2, 11, 12] in the human genome. Therefore, in order to comprehensively compare CGI features of the two reference genomes, we calculated not only all the density-defined and position-defined CGIs for the T2T-YAO-hp and T2T-CHM13 genomes but also the density-defined and position-defined CGIs related to divergent sequences of the two genomes (Supplementary **Figure S8**).

#### Density-defined CGI

Based on the GGF standard of CGIs [36], we computed the sequence segments that meet Equations. 1–3 as density-defined CGIs by the CpGPlot algorithm [42].

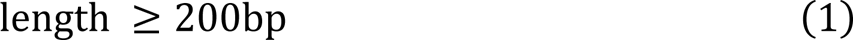

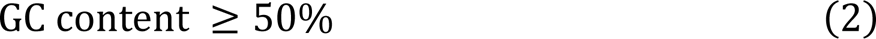

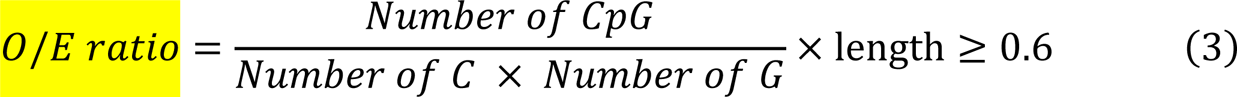

where length is the nucleotide number of the analysed gene sequence region, GC content is the proportion of G+C in the analysed gene sequence region, and O/E ratio represents the ratio of observed CpG to expected CpG.

#### Position-defined CGI

We first calculated the CpG clusters in the genome by Equation 4, and then considered these CpG clusters with small p values [26] (Equation 5) as position-defined CGIs.

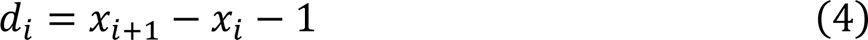

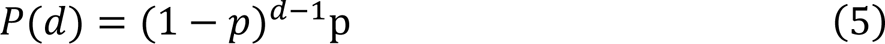

where *x* represents the coordinates of CpG and *i* represents the index of CpG. P(d) represents the probability of finding a distance d between neighboring CpGs. p corresponds to the probability of CpGs in the sequence. Since the default value of d in the CpGcluster algorithm is 50 and our previous study found that LAUPs (lineage-associated underrepresented permutations) are strongly associated with CGIs, the length of the shortest LAUPs of multiple species ranges from 10 to 14 [19], and the length of 25 bp is the statistical proportionality demarcation point for all CpGs and core-promoter-associated CpG regions in the human genome [2], we computed CGIs with d = 50 bp, 25 bp, and 12 bp (the average of 10 and 14).

### Genome-wide methylation analysis

#### WGBS data mapping

We applied gemBS [43], a large-scale WGBS data analysis method, to carry out methylation analysis for the two representative tissues (ovary and testis) by setting T2T-YAO-hp and T2T-CHM13 as references. The details of the procedure are described in Supplementary **Figure S9**.

#### CGI methylation ratio computing

After obtaining the CGI prediction for the reference genome and the methylation rate for each CpG site, Equation 6 calculates the methylation ratio for each CGI.

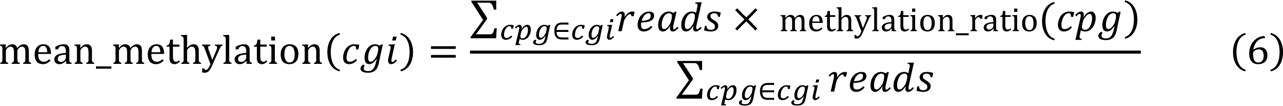

where *reads* represent the sequencing depth of WGBS for each CpG site in the CGI, and methylation_ratio(*cpg*) represents the methylation ratio of each CpG site derived from the gemBS parallelization algorithm.

#### Methylation level classification

According to the previous studies [2, 44], Equation 7 was used to classify the methylation ratio for each CpG site and CGI into three methylation levels, “hyper-methylated”, “hypo-methylated”, and “intermediate methylated”.

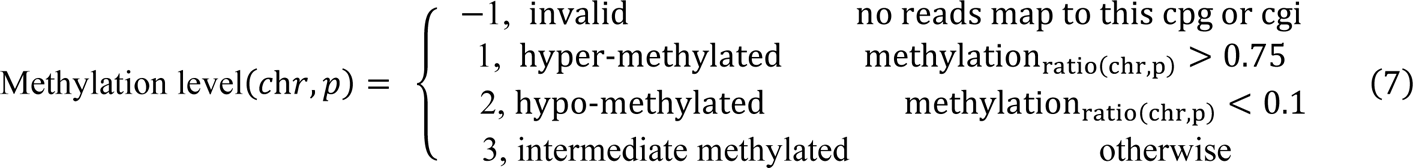

where chr and p represent the chromosome and the coordinates of the CpG site, respectively.

### CRediT author statement

**Ming Xiao:** Conceptualization, Methodology, Software, Validation, Investigation, Data curation, Writing - original draft, Writing - review & editing, Visualization, Supervision, Project administration, Funding acquisition. **Rui Wei:** Conceptualization, Methodology, Software, Validation, Investigation, Data curation, Writing - original draft, Writing - review & editing, Visualization, Project administration. **Jun Yu:** Conceptualization, Writing - review & editing, Visualization, Supervision, Project administration. **Chujie Gao:** Formal analysis, Investigation, Resources, Data curation. **Fengyi Yang:** Formal analysis, Investigation, Resources, Data curation. **Le Zhang:** Conceptualization, Writing - review & editing, Visualization, Supervision, Project administration, Funding acquisition. All authors have read and approved the final manuscript.

## Competing interests

The authors have declared no competing interests.

## Acknowledgments

This work was supported by grants from the National Natural Science Foundation of China [Grant No. 62372316], National Science and Technology Major Project (Nos. 2021YFF1201200 and 2018ZX10201002, China), China Postdoctoral Science Foundation (2020M673221, China), Fundamental Research Funds for the Central Universities (2020SCU12056, China), Sichuan Science and Technology Program (2022YFS0048), Chongqing Technology Innovation and Application Development Project (CSTB2022TIAD-KPX0067), China. We thank Dr. Yu Kang from Beijing Institute of Genomics, Chinese Academy of Sciences and China National Center for Bioinformation for her assistance in data analysis.

## ORCID

ORCID 0000-0001-8608-5903 (Ming Xiao)

ORCID 0009-0006-8597-5924 (Rui Wei)

ORCID 0000-0002-2702-055X (Jun Yu)

ORCID 0009-0000-3797-7040 (Chujie Gao)

ORCID 0009-0008-4105-030X (Fengyi Yang)

ORCID 0000-0002-3708-1727 (Le Zhang)

## Notes

### Summary of Updates

We've enhanced the paper by refining the language and improving the figures and tables, making it more precise and impactful.

