## Supplementary figures and images for "CpG Island Definition and Methylation Mapping of the T2T-YAO Genome"

### Supplementary Figure S5

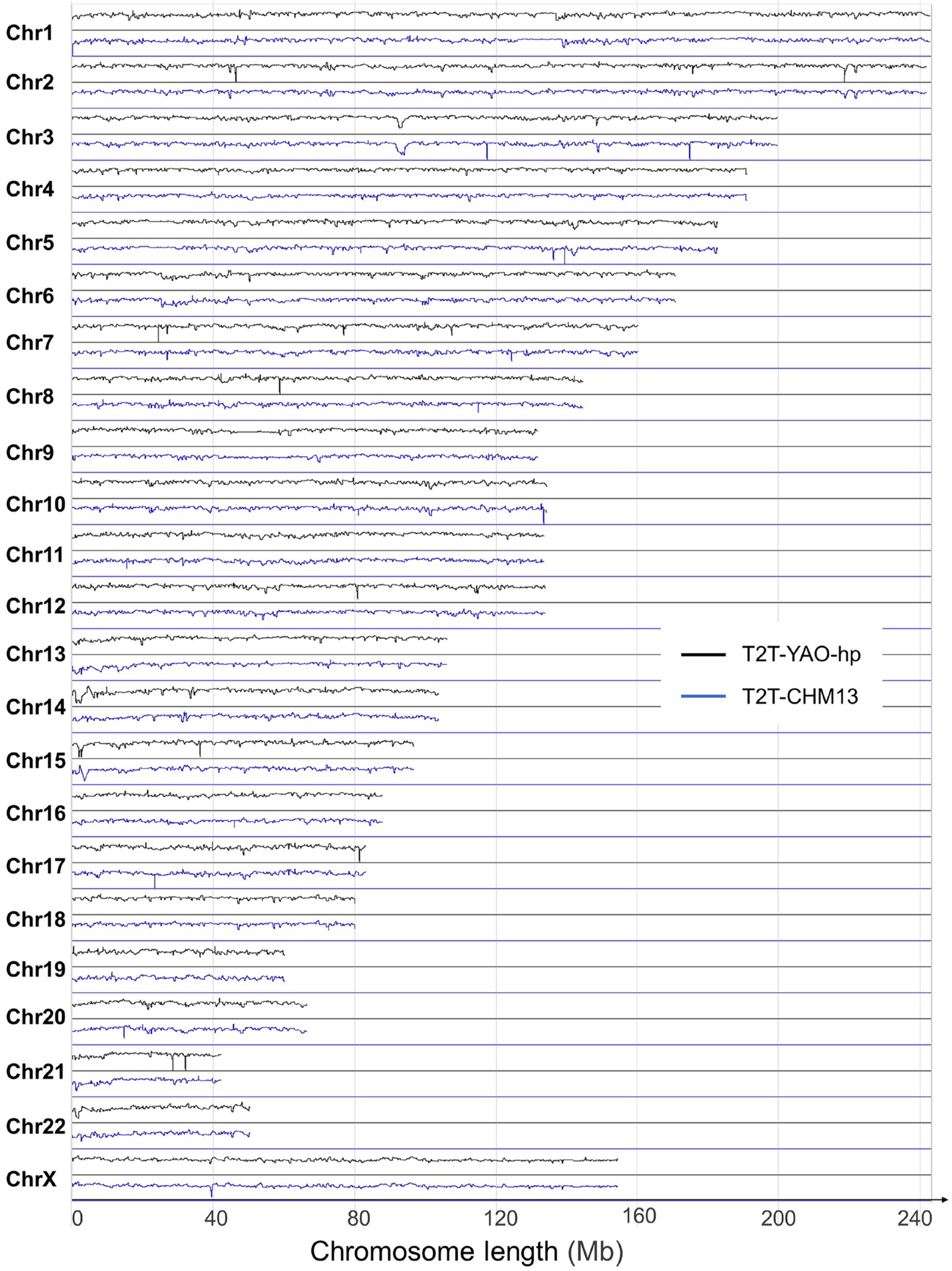

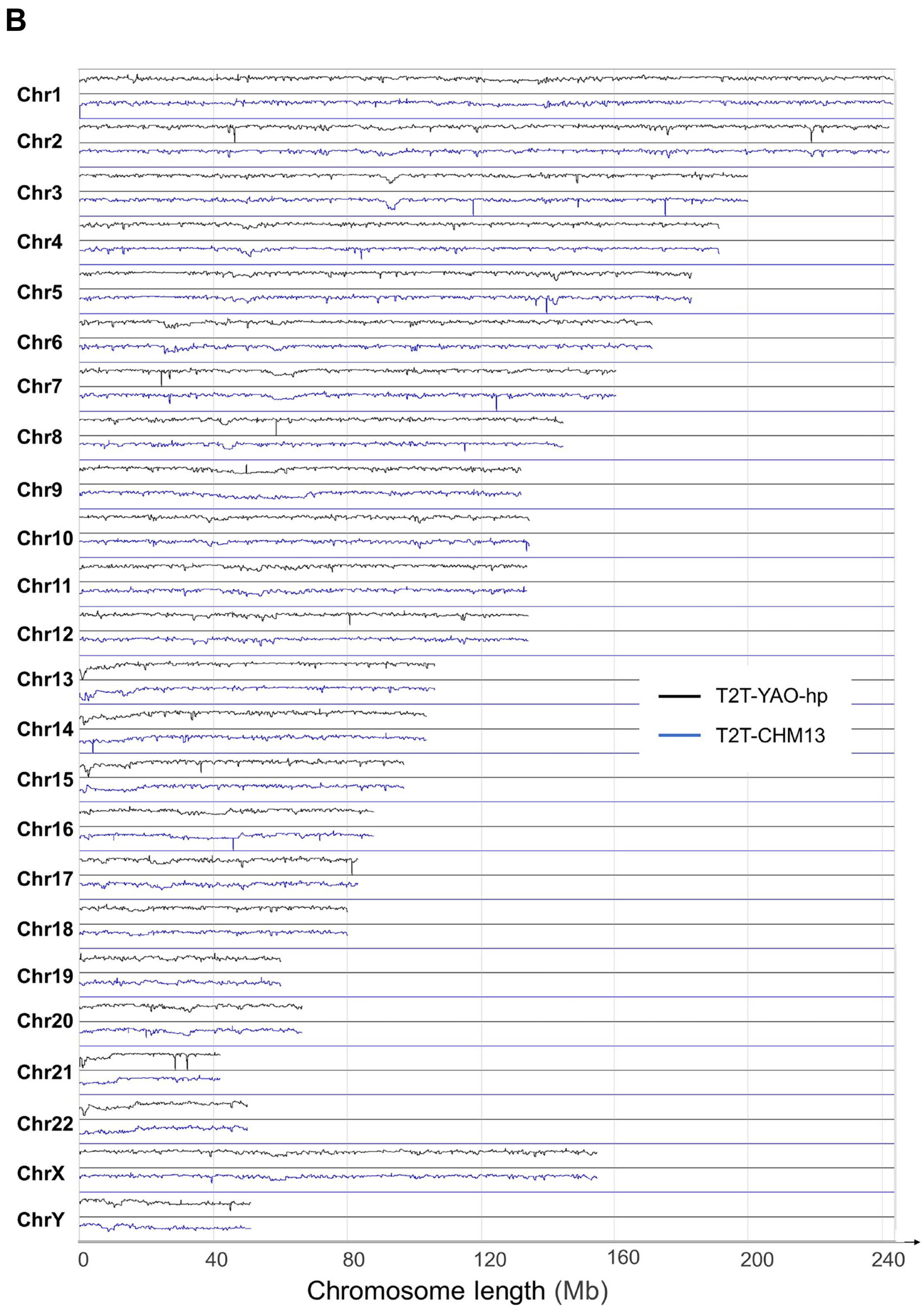

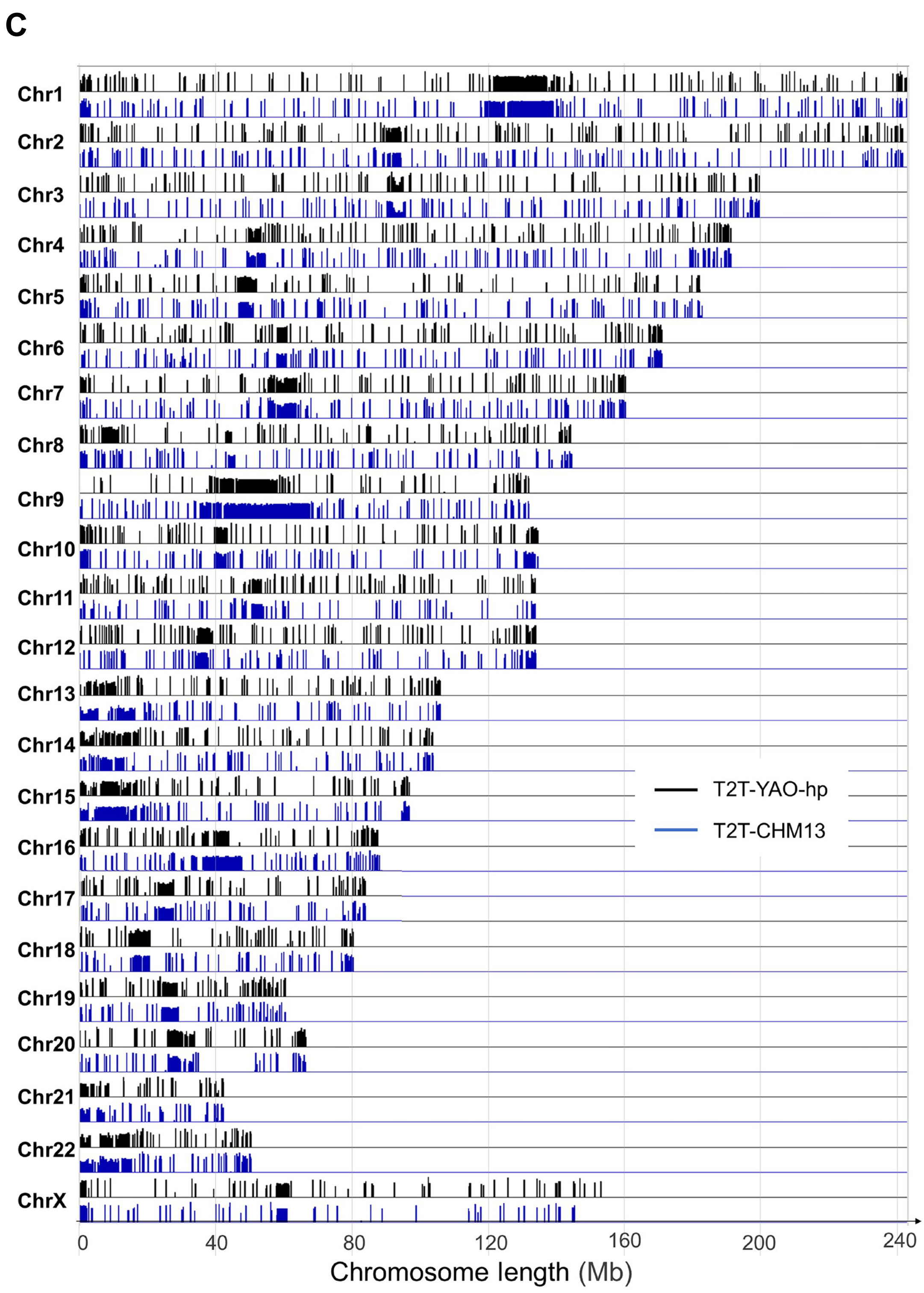

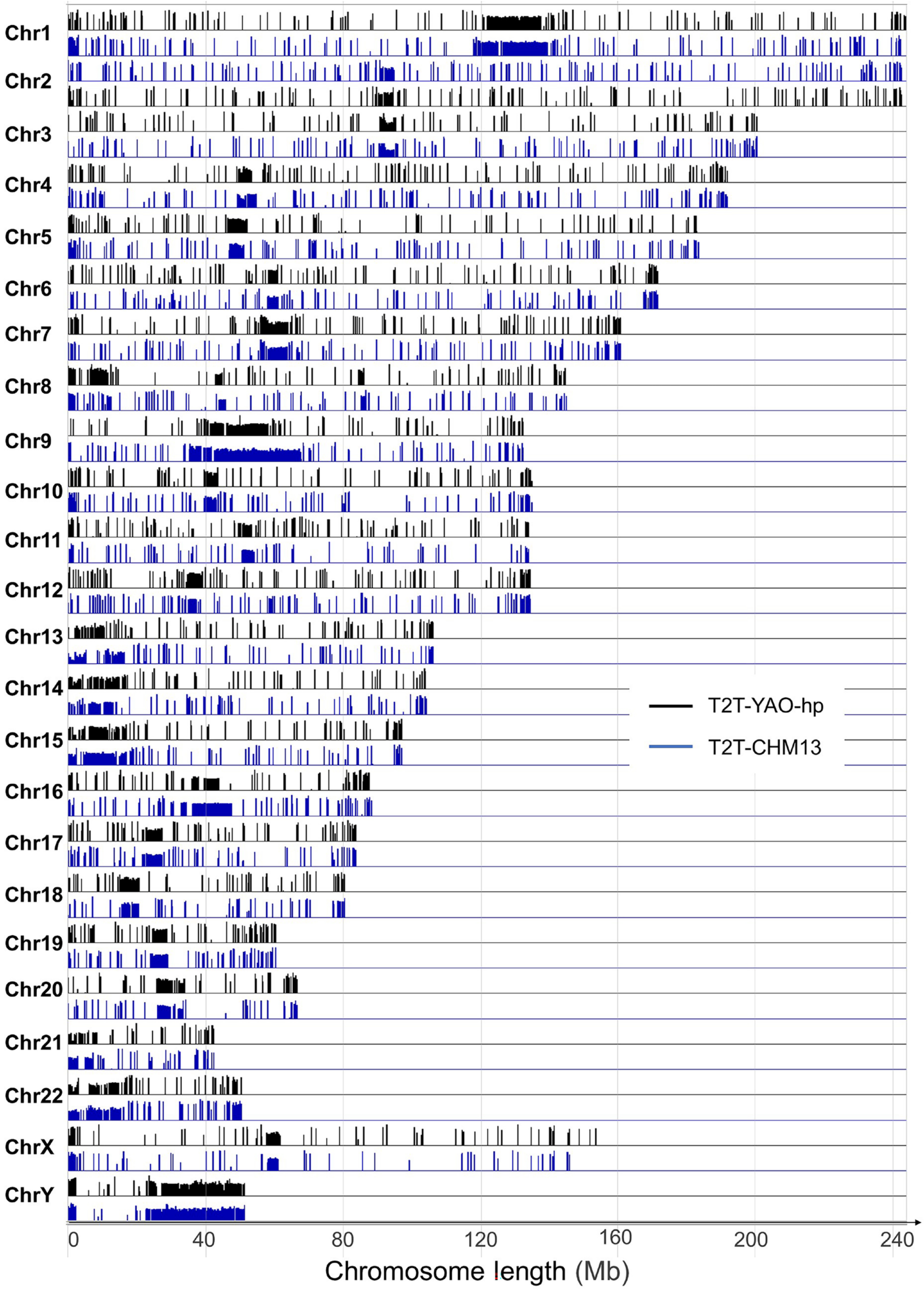
