## Supplementary Table S6 for "CpG Island Definition and Methylation Mapping of the T2T-YAO Genome"

**Table S6 Differential statistics of the distribution of CGI features between the two genomes**

|  | **CGI length** | **GC content** | **O/E ratio** | **CpG density** |
| --- | --- | --- | --- | --- |
| All density-defined CGIs | 0.0405 | 0.0191 | 0.0196 | 0.0213 |
| All position-defined CGIs | 0.0610 | 0.1067 | 0.0875 | 0.1561 |
| Divergent sequences associated Density-defined CGIs | 0.3188 | 0.1877 | 0.1676 | 0.1439 |
| Divergent sequences associated position-defined CGIs | 0.1237 | 0.2691 | 0.2594 | 0.3462 |

*Note:* In this table, we counted the sum of the pairwise differences of CGI features distribution between the two genomes (Figure 5). As can be observed from the table, the difference values of divergent sequence-associated CGIs are all larger than that of all CGIs (*e.g.*, 0.1877 *vs.* 0.0191 and 0.2691 *vs.* 0.1067 for GC content). This suggests that the difference in divergent sequences associated CGIs is more significant than all CGIs between the two genomes.
