## Supplementary information for "CpG Island Definition and Methylation Mapping of the T2T-YAO Genome"

**Supplementary material**

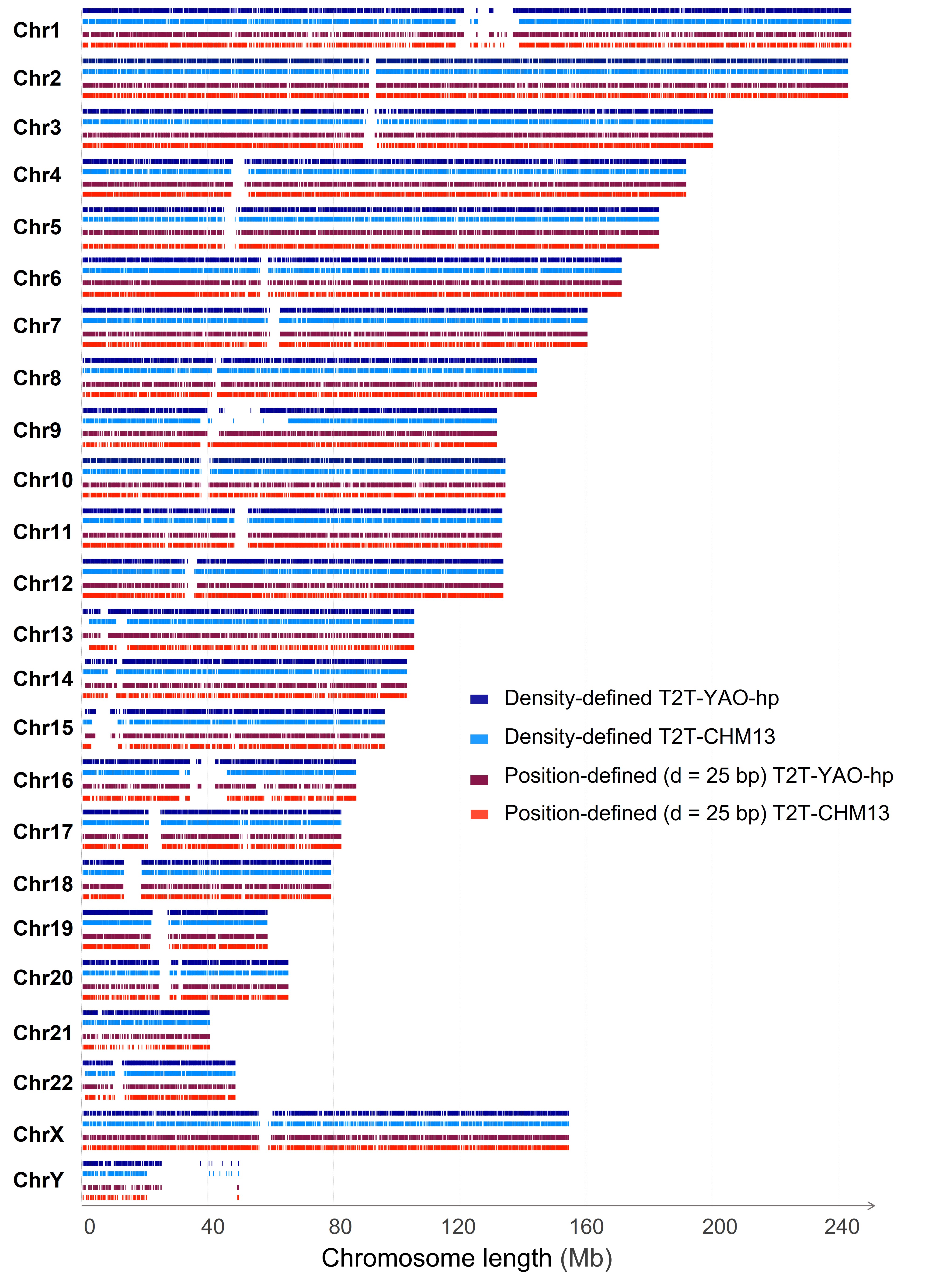

**Supplementary Figure S1 Comparison of CGI distribution on each chromosome in the T2T-YAO-hp and T2T-CHM13 genomes**

Blue and red represent density-defined CGI and position-defined CGI (d = 25 bp), respectively. Dark and light colors are used to distinguish T2T-YAO-hp and T2T-CHM13.

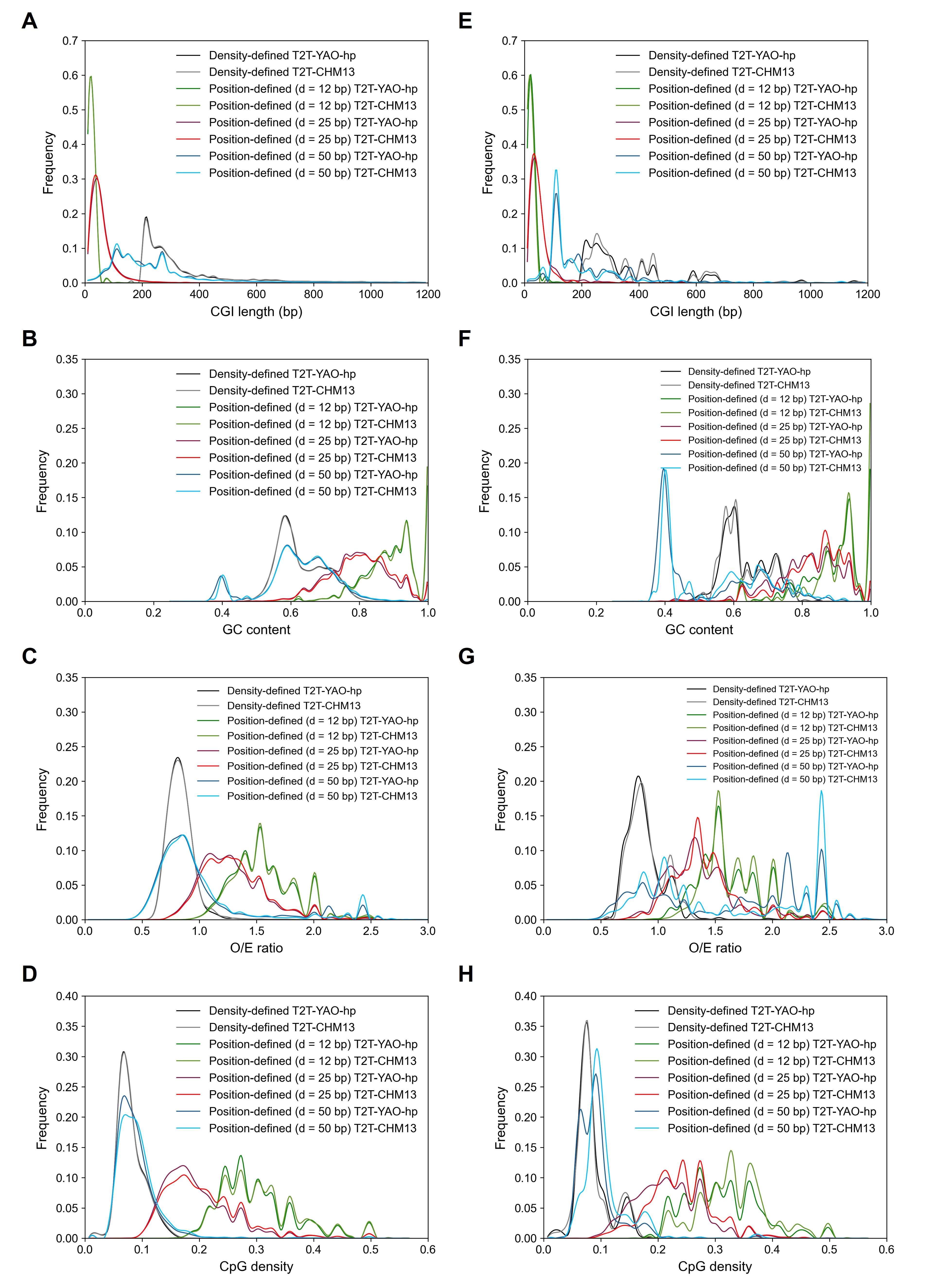

**Supplementary Figure S2 Comparison of CGI features between the T2T-CHM13 and T2T-YAO-hp genomes (including position-defined with d = 12 bp and d = 50 bp)**

**A.** Length of all CGIs; **B.** GC content of all CGIs; **C.** O/E ratio of all CGIs; **D.** CpG density of all CGIs; **E.** Length of divergent sequences CGIs; **F.** GC content of divergent sequences CGIs; **G.** O/E ratio of divergent sequences CGIs; **H.** CpG density of divergent sequences CGIs.

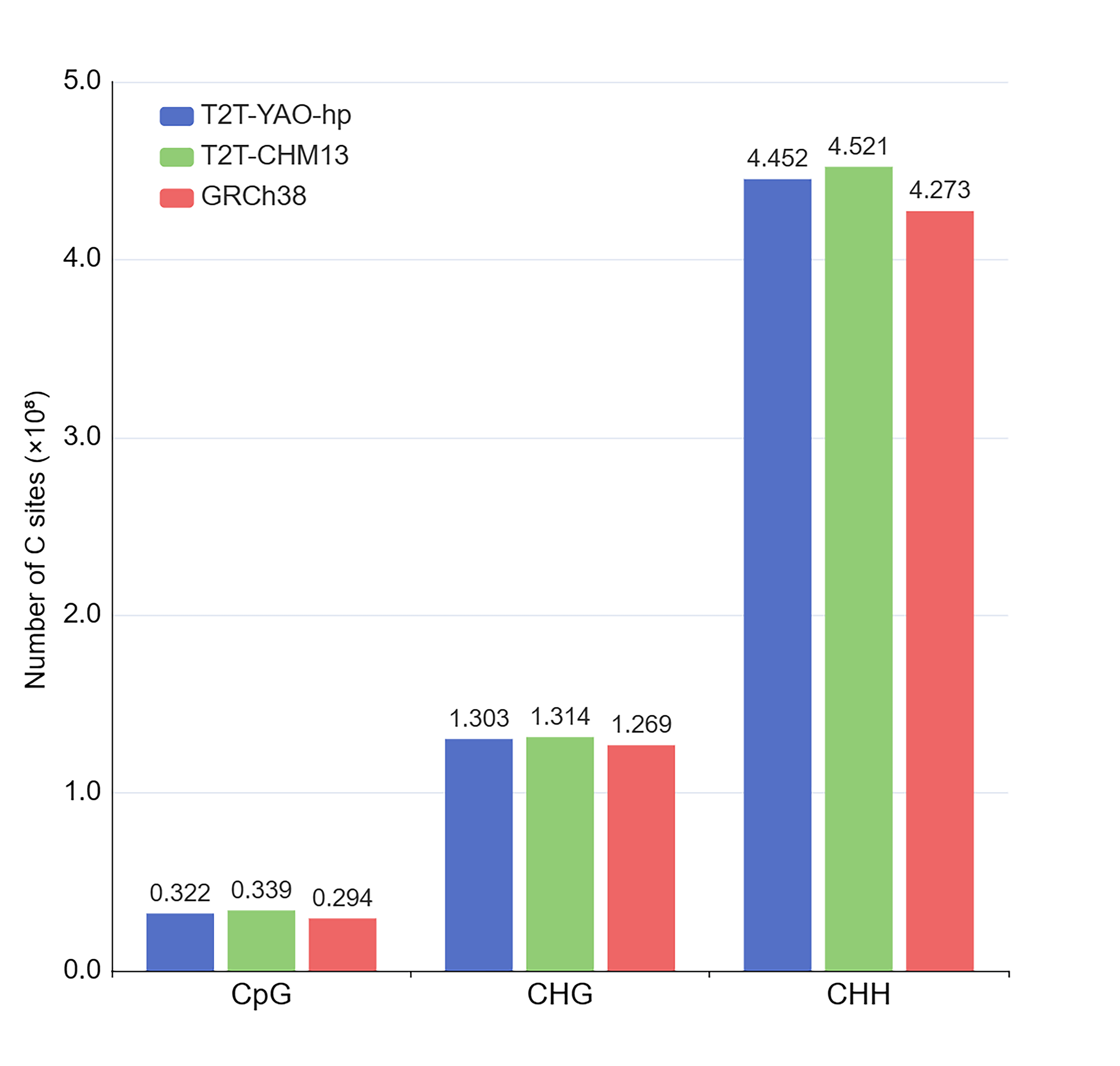

**Supplementary Figure S3 Comparison of C sites between the T2T-YAO-hp, T2T-CHM13 and GRCh38 genomes**

C sites were categorized into three different contexts: CpG, CHG, and CHH, where H represents A, C, or T.

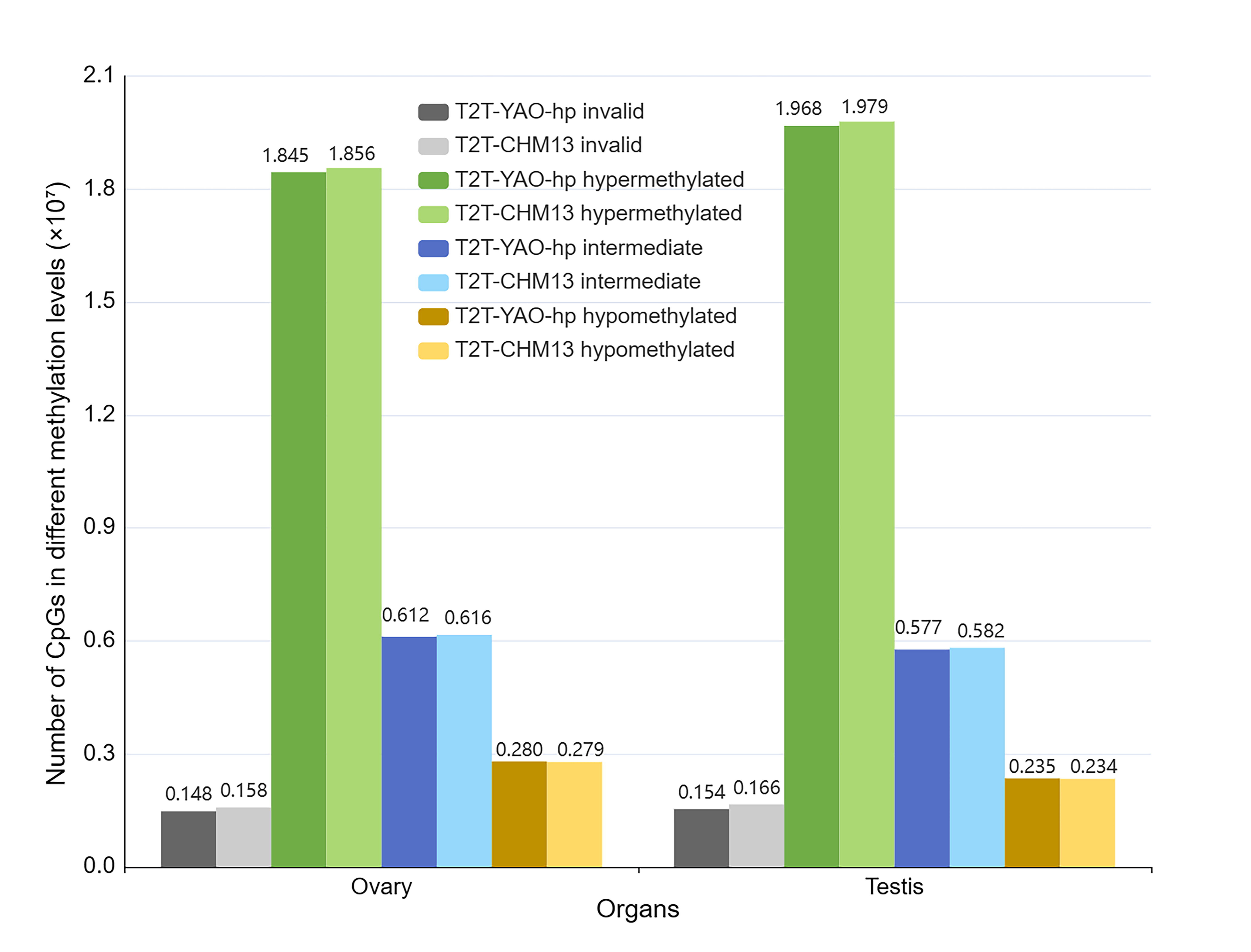

**Supplementary Figure S4 CpG methylation level of similar sequences between the T2T-YAO-hp and T2T-CHM13 genomes**

Green, blue, and orange represent the number of hyper-methylated, intermediate methylated and hypo-methylated, respectively; gray represents the number of CpG sites that failed to be mapped by WGBS data; dark and light colors are used to distinguish T2T-YAO-hp and T2T-CHM13.

**Supplementary Figure S5 Whole genome and the divergent sequences methylation profiles of the T2T-YAO-hp and T2T-CHM13 genomes**

**A.** Genome-wide methylation profiles of the ovary. **B.** Genome-wide methylation profiles of the testis. **C.** Divergent sequence methylation profiles of the ovary. **D.** Divergent sequence methylation profiles of the testis.

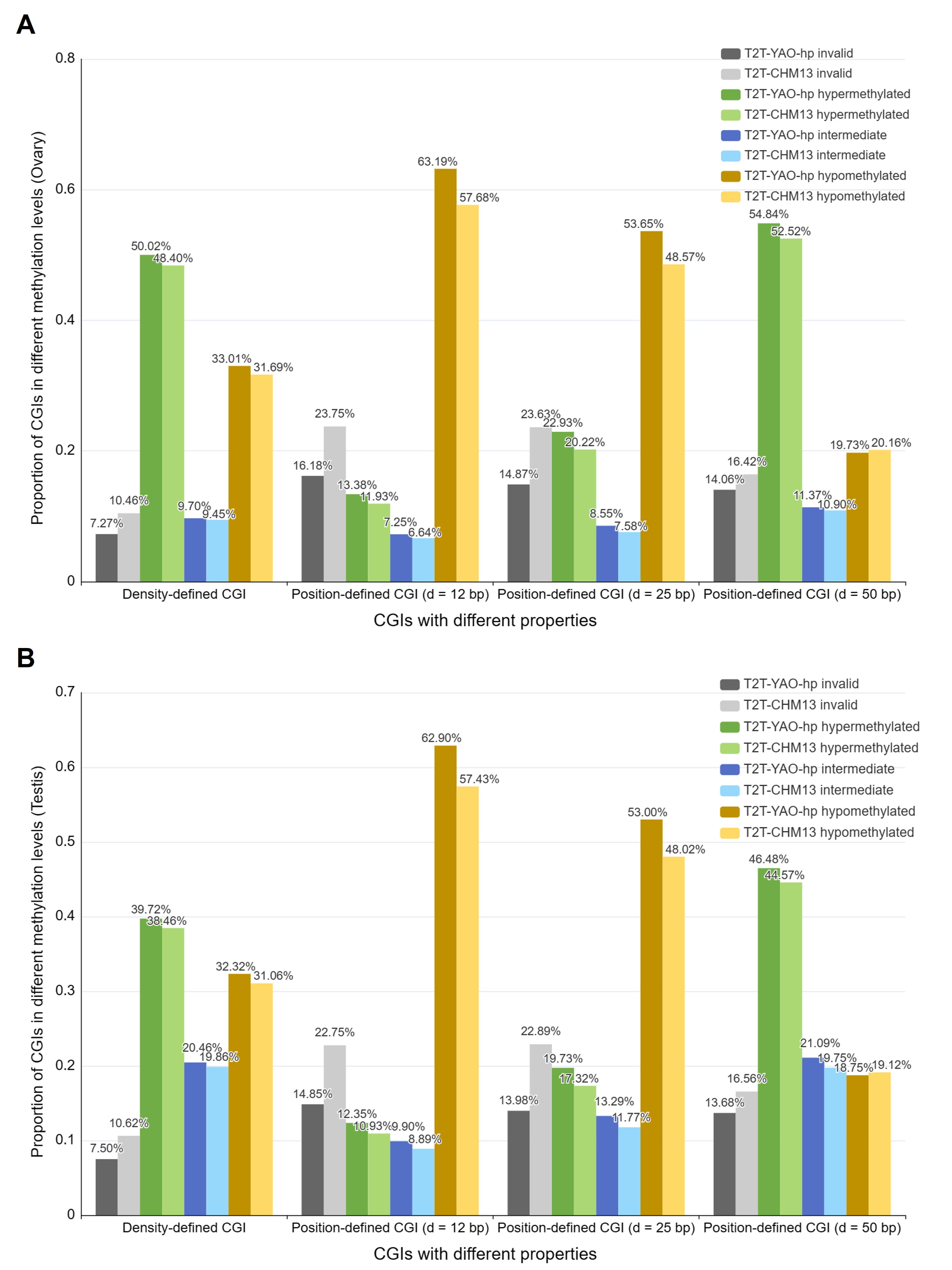

**Supplementary Figure S6 Comparison of CGl methylation between the T2T-YAO-hp and T2T-CHM13 genomes (including position-defined with d = 12 bp and d = 50 bp)**

**A.** CGI methylation level of two reference genomes using the WGBS data of Ovary. **B.** CGI methylation level of two reference genomes using the WGBS data of Testis. Here, green, blue, and orange represent the proportion of CGIs that are hyper-methylated, intermediate methylated, and hypo-methylated, respectively. Gray represents the proportion of CGIs where all CpGs failed to be mapped by WGBS data which named “invalid”. Darker and lighter colors are used to distinguish the T2T-YAO-hp and T2T-CHM13.

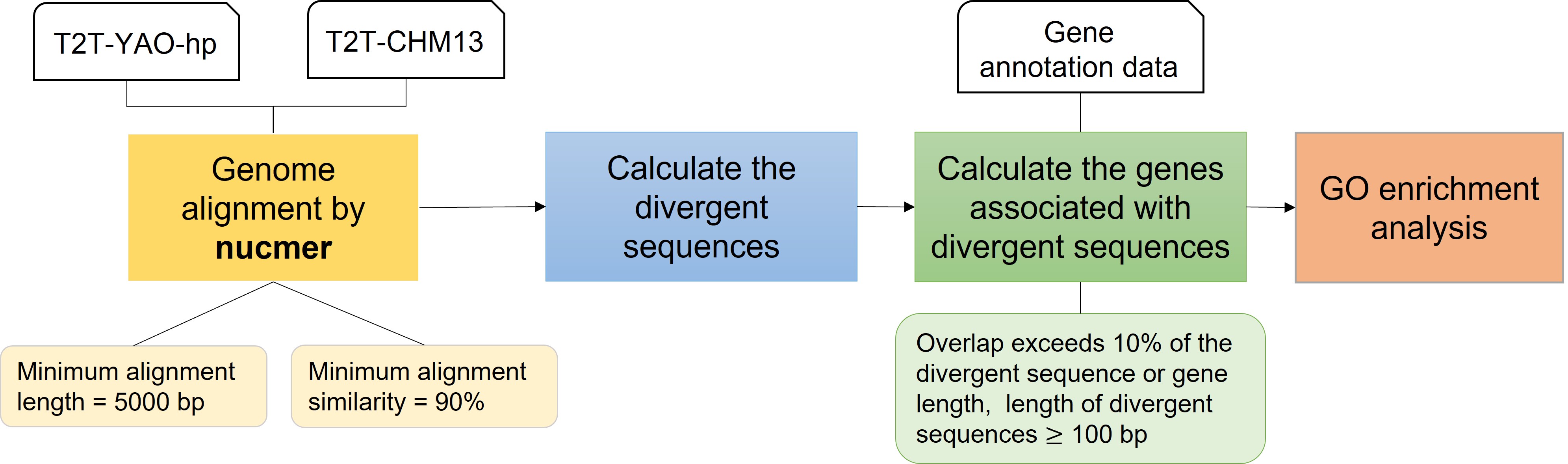

**Supplementary Figure S7 Flow chart of genome alignment and analysis**

First, we performed genome alignment of T2T-YAO-hp and T2T-CHM13 using nucmer to calculate the similar and divergent sequences of the two genomes. Next, the divergent sequence related genes were calculated and they were analyzed for GO enrichment.

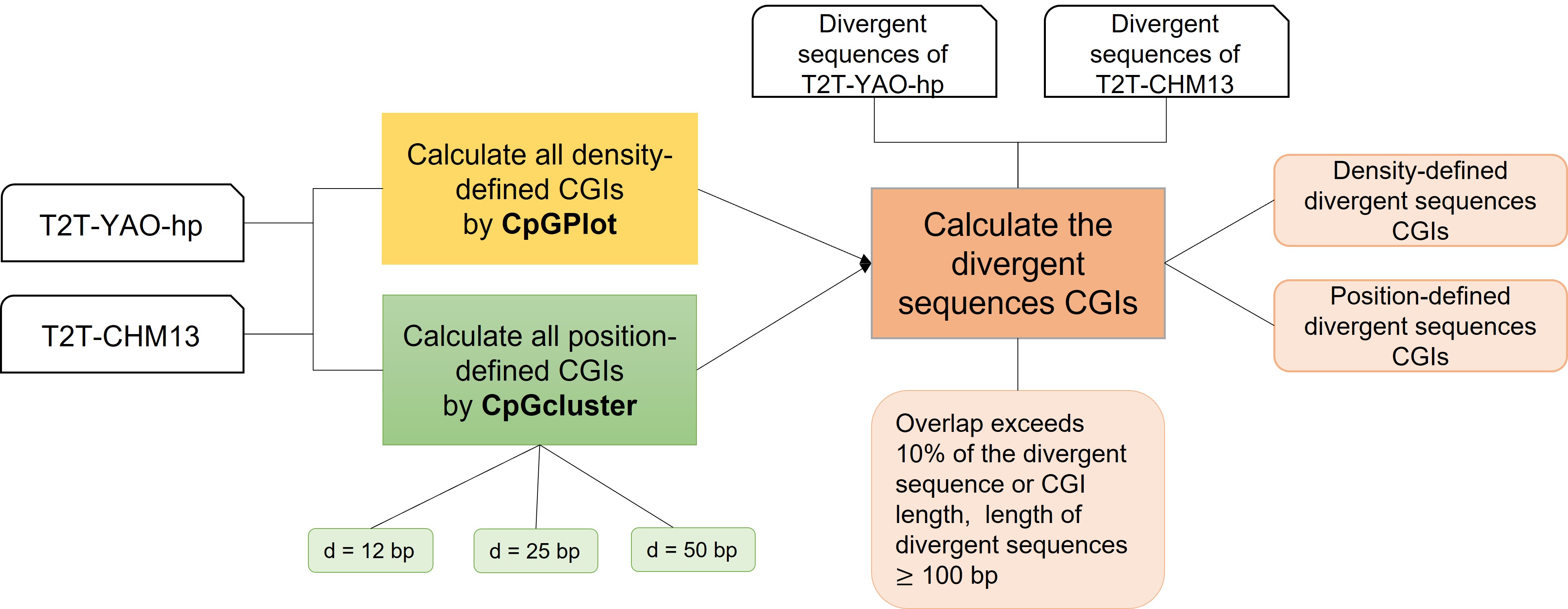

**Supplementary Figure S8 CGI detection in whole-genome and divergent sequences of the T2T-YAO-hp and T2T-CHM13 genomes**

First, we calculated density-defined CGIs and position-defined CGIs for T2T-YAO-hp and T2T-CHM13, respectively. Next, divergent sequence CGIs were calculated based on the degree of overlap of these CGIs with the divergent sequences.

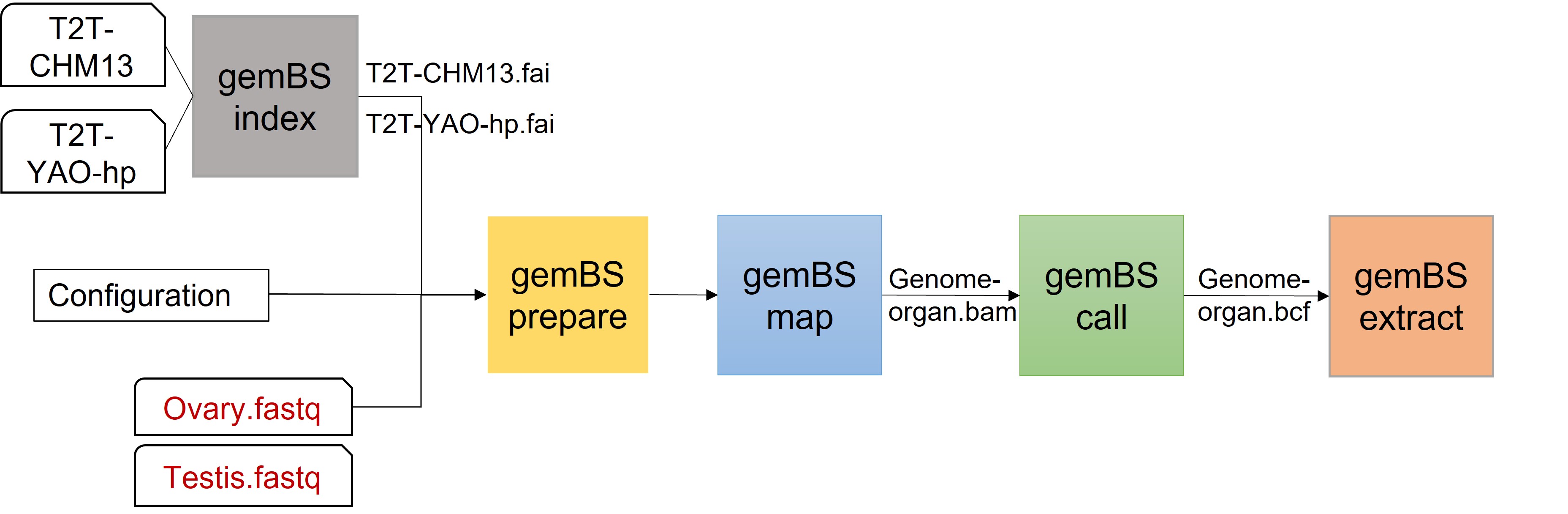

**Supplementary Figure S9 Flow chart of WGBS data mapping and calculation**

**Supplementary Table S1 Similar sequences between the T2T-YAO-hp and T2T-CHM13 genomes**

**Supplementary Table S2 Divergent sequences between the T2T-YAO-hp and T2T-CHM13 genomes**

**Supplementary Table S3 Divergent-sequence-associated genes**

**Supplementary Table S4 All CGIs for the T2T-YAO-hp and T2T-CHM13 genomes**

**Supplementary Table S5 All divergent-sequence-associated CGIs for the T2T-YAO-hp and T2T-CHM13 genomes**

**Supplementary Table S6 Differential statistics of the distribution of CGI features between the T2T-YAO-hp and T2T-CHM13 genomes**
